# A Network Integration Approach for Drug-Target Interaction Prediction and Computational Drug Repositioning from Heterogeneous Information

**DOI:** 10.1101/100305

**Authors:** Yunan Luo, Xinbin Zhao, Jingtian Zhou, Jinglin Yang, Yanqing Zhang, Wenhua Kuang, Jian Peng, Ligong Chen, Jianyang Zeng

**Affiliations:** Institute for Interdisciplinary Information Sciences, Tsinghua University, Beijing, China; School of Pharmaceutical Sciences, Tsinghua University, Beijing, China; Department of Computer Science, University of Illinois at Urbana-Champaign, Urbana, IL, USA

## Abstract

The emergence of large-scale genomic, chemical and pharmacological data provides new opportunities for drug discovery and repositioning. Systematic integration of these heterogeneous data not only serves as a promising tool for identifying new drug-target interactions (DTIs), which is an important step in drug development, but also provides a more complete understanding of the molecular mechanisms of drug action. In this work, we integrate diverse drug-related information, including drugs, proteins, diseases and side-effects, together with their interactions, associations or similarities, to construct a heterogeneous network with 12,015 nodes and 1,895,445 edges. We then develop a new computational pipeline, called DTINet, to predict novel drug-target interactions from the constructed heterogeneous network. Specifically, DTINet focuses on learning a low-dimensional vector representation of features for each node, which accurately explains the topological properties of individual nodes in the heterogeneous network, and then predicts the likelihood of a new DTI based on these representations via a vector space projection scheme. DTINet achieves substantial performance improvement over other state-of-the-art methods for DTI prediction. Moreover, we have experimentally validated the novel interactions between three drugs and the cyclooxygenase (COX) protein family predicted by DTINet, and demonstrated the new potential applications of these identified COX inhibitors in preventing inflammatory diseases. These results indicate that DTINet can provide a practically useful tool for integrating heterogeneous information to predict new drug-target interactions and repurpose existing drugs. The source code of DTINet and the input heterogeneous network data can be downloaded from http://github.com/luoyunan/DTINet.

## 1 Introduction

Computational prediction of drug-target interactions (DTIs) has become an important step in the drug discovery or repositioning process, aiming to identify putative new drugs or novel targets for existing drugs. Compared to *in vivo* or biochemical experimental methods for identifying new DTIs, which can be extremely costly and time-consuming [73], in *silico* or computational approaches can efficiently identify potential DTI candidates for guiding in vivo validation, and thus significantly reduce the time and cost required for drug discovery or repositioning. Traditional computational methods mainly depend on two strategies, including the molecular docking-based approaches [11, 17, 42] and the ligand-based approaches [29, 30]. However, the performance of molecular docking is limited when the 3D structures of target proteins are not available, while the ligand-based approaches often lead to poor prediction results when a target has only a small number of known binding ligands.

In the past decade, much effort has been devoted to developing the machine learning based approaches for computational DTI prediction. A key idea behind these methods is the “guilt-by-association” assumption, that is, similar drugs may share similar targets and vice versa. Based on this intuition, the DTI prediction problem is often formulated as a binary classification task, which aims to predict whether a drug-target interaction is present or not. A straightforward classification based approach is to consider known DTIs as labels and incorporate chemical structures of drugs and primary sequences of targets as input features (or kernels). Most existing prediction methods mainly focus on exploiting information from homogeneous networks. For example, Bleakley and Yamanishi [5] applied a support vector machine (SVM) framework to predict DTIs based on a bipartite local model (BLM). Mei et al. [39] extended this framework by combining BLM with a neighbor-based interaction-profile inferring (NII) procedure (called BLMNII), which is able to learn the DTI features from neighbors and predict interactions for new drug or target candidates. Xia et al. [76] proposed a semi-supervised learning method for DTI prediction, called NetLapRLS, which applies Laplacian regularized least square and incorporates both similarity and interaction kernels into the prediction framework. van Laarhoven et al. introduced a Gaussian interaction profile (GIP) kernel based approach coupled with regularized least square (RLS) for DTI prediction [66, 65]. Rather than regarding a drug-target interaction as a binary indicator, Wang and Zeng [71] proposed a restricted Boltzmann machine (RBM) model to predict different types of DTIs (e.g., activation and inhibition) on a multidimensional network.

In addition to chemical and genomic data, previous works have incorporated pharmacological or phenotypic information, such as side-effect [7, 41], transcriptional response data [24], drug-disease associations [70], public gene expression data [18, 53] and functional data [77] for DTI prediction. Heterogeneous data sources provide diverse information and a multi-view perspective for predicting novel DTIs. For instance, the therapeutic effects of drugs on diseases can generally reflect their binding activities to the targets (proteins) that are related to these diseases and thus can also contribute to DTI prediction. Therefore, incorporating heterogeneous data sources, e.g., drug-disease associations, can potentially boost the accuracy of DTI prediction and provide new insights into drug repositioning. Despite the current availability of heterogeneous data, most existing methods for DTI prediction are limited to only homogeneous networks or a bipartite DTI models, and cannot be directly extended to take into account heterogeneous node or topological information and complex relations among different data sources.

Recently, several computational strategies have been introduced to integrate heterogeneous data sources to predict DTIs. A network-based approach for this purpose is to fuse heterogeneous information through a network diffusion process, and directly use the obtained diffusion distributions to derive the prediction scores of DTIs [70, 10]. A meta-path based approach has also been proposed to extract the semantic features of DTIs from heterogeneous networks [20]. A collaborative matrix factorization has been developed to project the heterogeneous networks into a common feature space, which enables one to use the aforementioned homogeneous network based methods to predict new DTIs from the resulting single integrated network [79]. However, these approaches generally fail to provide satisfactory integration paradigms. First, directly using the diffusion states as the features or prediction scores may easily suffer from the bias induced by the noise and high-dimensionality of biological data and thus possibly lead to inaccurate DTI predictions. In addition, the hand-engineered features, such as meta-paths, often require expert knowledge and intensive effort in feature engineering, and hence prevent the prediction methods from being scaled to large-scale datasets. Moreover, collapsing multiple individual networks into a single network may cause substantial loss of network-specific information, since edges from multiple data sources are mixed without distinction in such an integrated network.

In this paper, we present DTINet, a novel network integration pipeline for DTI prediction. DTINet not only integrates diverse information from heterogeneous data sources (e.g., drugs, proteins, diseases and side-effects), but also copes with the noisy, incomplete and high-dimensional nature of large-scale biological data by learning low-dimensional but informative vector representations of features for both drugs and proteins. The low-dimensional feature vectors learned by DTINet capture the context information of individual networks, as well as the topological properties of nodes (e.g., drugs or proteins) across multiple networks. Based on these low-dimensional feature vectors, DTINet then finds an optimal projection from drug space onto target space, which enables the prediction of new DTIs according to the geometric proximity of the mapped vectors in a unified space. We have demonstrated the integration capacity of DTINet by unifying multiple networks that are related to drugs and proteins, and shown that incorporating additional network information can significantly improve the prediction accuracy. In addition, through comprehensive tests, we have demonstrated that DTINet can achieve substantial performance improvement over other state-of-the-art prediction methods. Furthermore, we have experimentally validated the new interactions predicted by DTINet between three drugs and the cyclooxygenase (COX) protein family that have not been reported in the literature (to the best of our knowledge) and have demonstrated the potential novel applications of these drugs in preventing inflammatory diseases.

With superior prediction performance, DTINet offers a practically useful tool to predict unknown drug-target interactions from complex heterogeneous networks, which can provide new insights into drug discovery or repositioning and the understanding of mechanisms of drug action. Overall, we feature the following major advancements of DTINet: (i) the generalizability and scalability of integrating a variety of heterogeneous data sources; (ii) the ease of automated compact feature learning without any hand-engineered feature extraction or domain-specific expert knowledge; (iii) the ability of addressing the computational challenges in network integration arising from the high-dimensional, incomplete and noisy biological data; (iv) the high prediction accuracy and substantial performance improvement over previous methods; and (v) the novel predicted drug-target interactions that have been validated experimentally.

## 2 Methods

As an overview (Figure 1), DTINet integrates diverse information from heterogeneous network by first combining the network diffusion algorithm (random walk with restart, RWR [60]) and a dimensionality reduction scheme (diffusion component analysis, DCA [12]), to obtain informative, but low dimensional vector representations of nodes in the network. We call this process as *compact feature learning.* Intuitively, the low-dimensional feature vector obtained from this process encodes the relational properties (e.g., similarity), association information and topological context of each drug (or protein) node in the heterogeneous network. Next, DTINet finds the best projection from drug space onto protein space, such that the mapped feature vectors of drugs are geometrically close to their known interacting targets. After that, DTINet infers new interactions for a drug by ranking its target candidates according to their proximity to the projected feature vector of this drug. A key insight of our approach is that the drugs (or proteins) with similar topological properties in the heterogeneous network are more likely to be functionally correlated. For example, those drugs that are close in the directions of their feature vectors are more likely to act on the same target, and vice versa. This intuition allows us to predict unknown drug-target interactions by fully exploiting our previous knowledge about known drug-target interactions.

**Figure 1:**
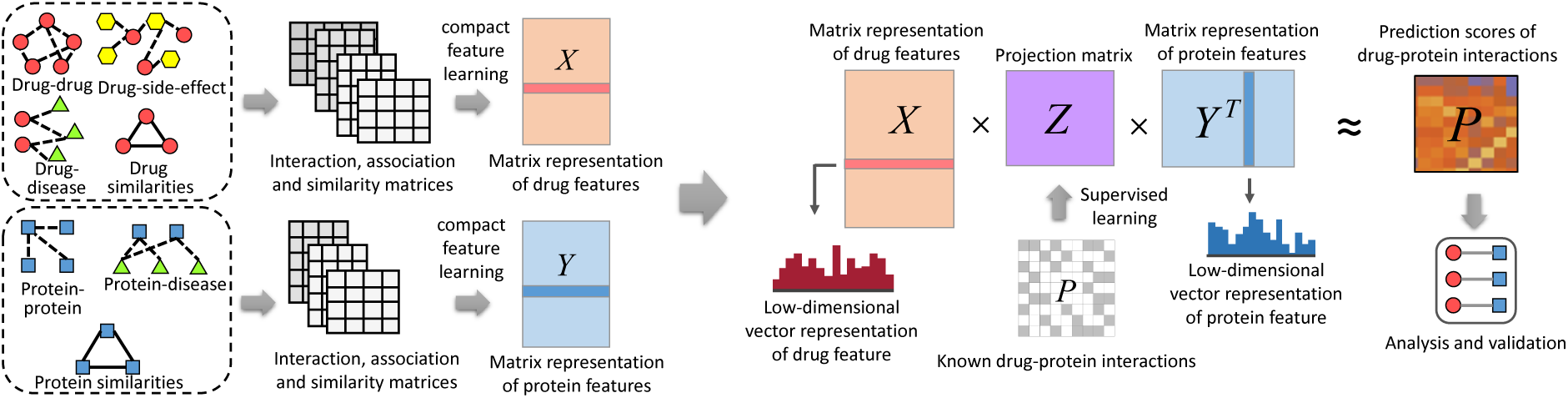
The flowchart of the DTINet pipeline. DTINet first integrates a variety of drug-related information sources to construct a heterogeneous network and applies a compact feature learning algorithm to obtain a low-dimensional vector representation of the features describing the topological properties for each node. DTINet then finds the best projection from drug space onto protein space, such that the projected feature vectors of drugs are geometrically close to the feature vectors of their known interacting proteins. After that, DTINet infers new interactions for a drug by sorting its target candidates based on their geometric proximity to the projected feature vector of this drug in the projected space. The predicted new drug-target interactions can be further analyzed and experimentally validated.

### 2.1 Compact feature learning for drugs and targets

#### Random walk with restart revisited

The first step of DTINet is to perform a network diffusion algorithm to jointly integrate heterogeneous information. Random walk with restart (RWR), a network diffusion algorithm, has been extensively applied to analyze the complex biological network data [8, 33, 44, 35, 10]. Different from conventional random walk methods, RWR introduces a pre-defined restart probability at the initial node for every iteration, which can take into consideration both local and global topological connectivity patterns within the network to fully exploit the underlying direct or indirect relations between nodes. Formally, let **A** denote the weighted adjacency matrix of a molecular interaction network with *n* drugs (or targets). We also define another matrix **B**, in which each element **B**_*i,j*_ describes the probability of a transition from node *i* to node *j*, that is, **B**_*i,j*_=**A**_*i,j*_/Σ _*j′*_ **A**_*i,j*_′. Next, let **s***_i_^t^* be an *n*-dimensional distribution vector in which each element stores the probability of a node being visited from node *i* after *t* iterations in the random walk process. Then RWR from node *i* can be defined as:

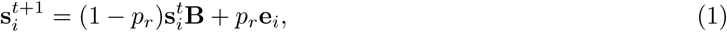

where **e**_*i*_ stands for an *n*-dimensional standard basis vector with **e**_*i*_(*i*) = 1 and **e**_*i*_ (*j*) = 0, ∀_*j*_ ≠ *i*, and *p_r_* stands for the pre-defined restart probability, which actually controls the relative influence between local and global topological information in the diffusion process, with higher values emphasizing more on the local structures in the network. At some fixed point of the iterating process, we can obtain a stationary distribution **s**_*i*_^∞^ of RWR, which we refer to as the “diffusion state” **s**_*i*_ for node *i* (i.e., **s**_*i*_ = **s**_*i*_^∞^), being consistent with the notation of previous work [12]. Intuitively, the *j*th element of diffusion state, denoted by **s**_*ij*_, represents the probability of RWR starting node *i* and ending up at node *j* in equilibrium. When two nodes have similar diffusion states, it generally implies that they have similar positions with respect to other nodes in the network, and thus probably share similar functions. This insight provides the basis for several previous works [70, 10] that aim to predict unknown DTIs based on diffusion states.

#### The dimensionality reduction framework

Although the diffusion states resulting from the aforementioned RWR process characterizes the underlying topological context and inherent connection profiles of each drug or protein node in the network, they may not be entirely accurate, partially due to the low-quality and high-dimensionality of biological data. For example, a small number of missing or fake interactions in the network can significantly affect the results of the diffusion process [31]. Moreover, it is generally inconvenient to directly use the high dimensional diffusion states as topological features in prediction tasks, especially in our heterogeneous network-based prediction task. To address this issue, DTINet employs a dimensionality reduction scheme, called diffusion component analysis (DCA), to reduce the dimensionality of feature space and capture those important topological features from the diffusion states. The key idea of DCA is to obtain informative but low dimensional vector representations, which encode the connectivity relations and topological properties of each node in the heterogeneous network. Akin to principal component analysis (PCA), which seeks the intrinsic low-dimensional linear structure of the data to best explain the variance, DCA learns a low-dimensional vector representation for all nodes such that their connectivity patterns in the heterogeneous network are best interpreted. We give a brief description of the DCA framework below.

With the goal of denoise and dimensionality reduction, DCA approximates the obtained diffusion state distribution **s**_*i*_ with a multinomial logistic model parameterized by a low-dimensional vector representation whose dimensionality is much lower than that of the original *n*-dimensional vector representing the diffusion states. Specifically, the probability assigned to node *j* in the diffusion state of node *i* is now modeled as

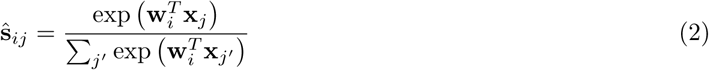

where ∀_*i*_, **x**_*i*_, **w**_*i*_ ∈ ℝ^*d*^ for *d* ≪ *n*. We refer to **w**_*i*_ as the *context feature* and **x**_*i*_ as the *node feature* for node *i*, both describing the topological properties of the network. If **x**_*i*_ and **w**_*j*_ point to a similar direction and thus have a large inner product, it is likely that node *j* is frequently visited in a random walk starting from node *i*. DCA takes a set of the observed diffusion states ***S*** = {**s**_1_,…, **s**_*n*_} as input and optimizes over **w** and **x** for all nodes, using the Kullback-Leibler (KL) divergence (also called relative entropy) to guide the optimization, that is,

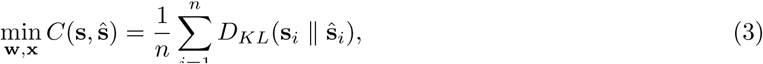

where *D_KL_*(· || ·) denotes the KL-divergence between two distributions. The DCA framework uses a standard quasi-Newton method L-BFGS [80] to solve this optimization problem.

#### Integration of heterogeneous network information

The above dimensionality reduction framework can be naturally extended to integrate multiple network data from heterogeneous sources. Given K similarity networks in a heterogeneous framework constructed from diverse information (Supplementary Information), DCA first performs RWR on individual networks separately and then obtains the network-specific diffusion states s*_i_^(k)^* for each node *i* in every network *k*. After that, it also constructs a multinomial logistic distribution to model the diffusion states:

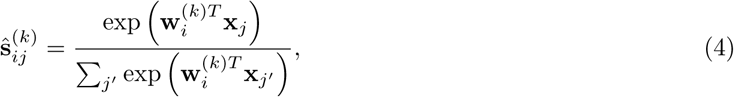

where each node *i* is assigned with a network-specific vector representation **w***_i_^(k)^*, which represents the context feature of node *i* in network *k*, and the node feature vectors x_*<u>i</u>*_ are allowed to be shared globally across all **K** networks. Finally, DCA optimizes the following objective function,

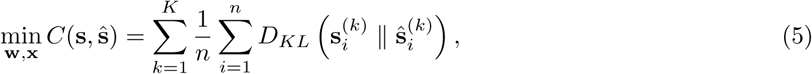

which can also be solved by the quasi-Newton L-BFGS method [80]. Although the divergence terms of individual networks are given equal weights in the above objective function, it is possible to weight them differently to emphasize the relative importance of individual networks. To make the DCA framework more scalable to large biological networks, DTINet employs an extended version of DCA, called clusDCA [69], which uses an alternative objective function that can be optimized efficiently based on singular value decomposition (SVD) (Supplementary Information).

### 2.2 Projection from drug space onto target space

We use the low-dimensional vector representations of both drug and protein features obtained from compact feature learning to predict new drug-target interactions. Based on the intuition that geometric proximity in the feature vector space may reflect the functional relevance, we apply a matrix completion approach [43] to obtain a projection matrix that maps the low-dimensional feature vectors from drug space onto protein space, such that the projected feature vectors of drugs are geometrically close to the vectors of their known interacting proteins. The insight behind this is that the drugs (or proteins) with similar topological properties in the heterogeneous network are more likely to be functionally correlated. For example, those drugs that are close in the directions of their feature vectors are more likely to share the same target, and vice versa. Based on this intuition, we can fully exploit previous knowledge of known DTIs to predict unknown DTIs.

Formally, we use **X** = [*x*_1_,…,*x_N_d__*]^*T*^, *x_i_* ∈ ℝ*^f_d_^, i* = 1, …,*N_d_* to denote the matrix representation of the drug features (i.e., each row *i* represents the corresponding feature vector of drug *i*), and **Y** = [*y*_1_,…,*y_N_t__*]^*T*^, *y_i_* ∈ ℝ*^f_t_^, i* = 1, …, *N_t_*, to denote the matrix representation of the protein features (i.e., each row *i* represents the corresponding feature vector of protien *i*), where *N_d_* and *N_t_* stand for the numbers of drugs and proteins, respectively (Figure 1). Let **P** be a drug-target interaction matrix, where each entry **P**_*ij*_ = 1 if drug *i* is known to interact with protein *j*, and **P**_*ij*_ = 0 otherwise. We set up a bilinear function to learn the projection matrix **Z** between drug space and target space to predict the unknown drug-target interactions in **P** (i.e., those zero-valued entries). In particular, the bilinear function is formulated as **XZY**^*T*^ ≈ **P**, where **P** ∈ ℝ*^N_d_ × N_t_^* stands for the known drug-target interaction matrix, **X** ∈ ℝ*^N_d_ × f_d_^*, **Y** ∈ ℝ*^N_t_ × f_t_^* are obtained from the compact feature learning stage (i.e., the network diffusion and dimensionality reduction processes), and **Z** ∈ ℝ*^f_d_ × f_t_^* is the projection matrix to be learned. We then use the formula below to measure the likelihood of the pairwise interaction score between drug *i* and protein *j*:

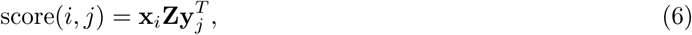

where a larger score(*i*, *j*) suggests that drug *i* is more likely to interact with protein *j*.

Although the projection matrix **Z** is of dimension *f_d_* × *f_t_*, there typically exist significant correlations between those feature vectors of drugs or proteins that are geometrically close in space, which can thus greatly reduce the number of effective parameters required in **Z** to model drug-target interactions. To take into account this issue, we impose a low-rank constraint on **Z** to learn only a small number of latent factors, by considering a low-rank decomposition of the form **Z** = **GH**^*T*^, where **G** ∈ ℝ*^f_d_ × f_k_^* and **H** ∈ ℝ*^f_t_ × f_k_^*. This low-rank constraint not only alleviates the overfitting problem but also computationally benefits the optimization process [78]. The optimization problem with such a low-rank constraint on the original projection matrix **Z** is NP-hard to solve. A standard relaxation of the low-rank constraint is to minimizing the trace norm (i.e., sum of singular values) of the matrix **Z** = **GH**^*T*^, which is equivalent to minimize 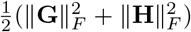. Therefore, factoring **Z** into **G** and **H** can be accomplished by solving the following optimization problem by alternating minimization [43]:

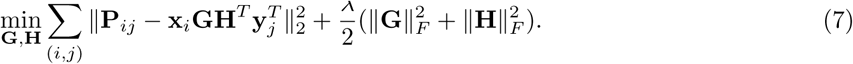

## 3 Results

Compiling various curated public drug-related databases, we constructed a heterogeneous network, which includes 12,015 nodes and 1,895,445 edges in total, for predicting missing drug-target interactions (Figure 2a). The heterogeneous network integrates four types of nodes (i.e., drugs, proteins, diseases and side-effects) and six types of edges (i.e., drug-protein interactions, drug-drug interactions, drug-disease associations, drug-side-effect associations, protein-disease associations and protein-protein interactions). We compared DTINet with four state-of-the-art methods of DTI prediction methods, including BLMNII [39], NetLapRLS [76], HNM [70] and CMF [79]. The area under the receiver operating characteristic curve (AUROC) and the area under the precision recall curve (AUPR) were used to assess the performance of each DTI prediction method. A detailed description of the dataset and experimental settings can be found in Supplementary Information.

**Figure 2:**
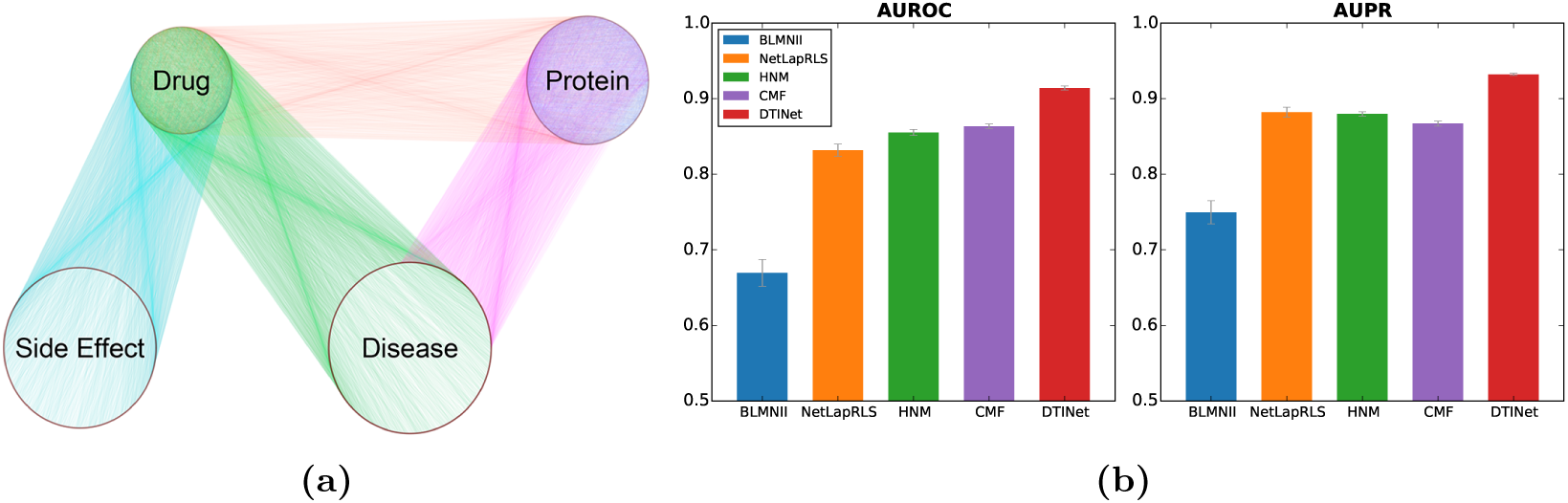
(a) Overview of the heterogeneous network constructed from diverse information to predict drug-target interactions. (b) DTINet outperforms other state-of-the-art methods for DTI prediction.

### 3.1 DTINet yields accurate drug-target interaction prediction

We first evaluated the prediction performance of DTINet using ten independent trials of ten-fold cross-validation procedure, in which a randomly chosen subset of 10% of the known interacting drug-target pairs and a matching number of randomly sampled non-interacting pairs were held out as the test set, and the remaining 90% known interactions and a matching number of randomly sampled non-interacting pairs were used to train the model. Our comparative results showed that DTINet consistently outperformed other existing methods in terms of both AUROC and AUPR scores (Figure 2b). For example, DTINet achieved an AUROC of 91.41% and an AUPR of 93.22%, which were 5.9% and 5.7% higher than the second best method, respectively. We found that DTINet had a clear improvement in prediction performance over the single network based approaches, i.e., BLMNII and NetLapRLS. Compared with other integrative methods, e.g., HNM, which predicts DTIs based on a modified version of random walk in a complete heterogeneous network, DTINet achieved 6.9% higher AUROC (85.51% for HNM) and 5.9% higher AUPR (87.98% for HNM), presumably due to the fact that HNM only uses the original diffusion states for prediction, which suffer from noise in the data and are not entirely accurate, while DTINet applies a novel dimensionality reduction on the diffusion states and thus is able to capture the underlying structural properties of the heterogeneous network.

To mimic a practical situation in which a drug-target interaction matrix is often sparsely labeled with only a few known DTIs, we also performed an additional ten-fold cross-validation test (Figure S3a), in which a much larger set of non-interacting drug-target pairs were included, such that the known drug-target interactions composed only 10% of the training or test dataset. This additional test showed that DTINet can still achieve decent performance and outperforms other prediction methods, even when only sparsely labeled DTIs are available to train the model. More specifically, DTINet achieved AUROC 91.52% and AUPR 77.72% on this skewed dataset, which were 6.0% and 21.5% higher than the second best baseline method, respectively. We also want to highlight the pronounced gap between DTINet and other prediction methods in terms of AUPR (138.7%, 21.5%, 38.4% and 64.4% higher than BLMNII, NetLapRLS, HNM and CMF, respectively). As studied in a previous work [15, 6, 66], AUROC is likely to be an overoptimistic metric to evaluate the performance of an algorithm in DTI prediction task, especially on highly-skewed data, while AUPR can provide a better assessment in this scenario. The noticeable performance improvement of DTINet in terms of AUPR over other prediction methods demonstrated its superior ability in predicting new drug-target interactions in the sparsely labeled networks.

The original collected dataset (Figure 2a, Table S1) may contain homologous proteins with high sequence identity scores, which raised a potential concern that DTINet’s good performance might result from easy predictions, in which the predicted targets had high sequence identity scores with the corresponding homologous proteins in training data. To investigate this issue, as in [71], we performed an additional test, in which we only kept those DTIs in which proteins had sequence identity scores < 40% in both training and test data. More precisely, for each group of proteins that had pairwise sequence identity scores > 40%, we only retained the protein that had the largest number of interacting drugs and removed all other proteins in that group. The removal of homologous proteins reduced the number of known DTIs from 1,923 to 1,332 in the dataset. We then assessed the performance of DTINet and other prediction methods again on this resulting dataset (Figure S2). We found that DTINet was robust to the removal of homologous proteins in training data, and still consistently outperformed other methods on such a dataset. For example, DTINet yielded AUROC 88.22% and AUPR 90.45%, which were significantly higher than the second best method. We also removed homologous proteins in the skewed dataset and compared the performance of DTINet to that of other baseline methods, and still observed superior performance of DTINet over others under this setting (Figure S3b). Taken together, these results demonstrated the superiority of DTINet in predicting novel DTIs, allowing us to find new DTIs that were not present in the original dataset (see the next section).

Our further comparative study showed that integrating multiple networks derived from the feature vectors of drugs or proteins by DTINet can greatly improve the prediction performance over individual single networks (Figure S4). Our comparison demonstrated that, even without multiple networks integration, DITNet still outperformed the state-of-the-art single network based method NetLapRLS on individual similarity networks. This result emphasised DTINet’s ability to fully exploit useful topological information from high-dimensional and noisy network data via a compact feature learning procedure, even only given a single network as input. In addition, we observed that DTINet achieved much better prediction performance than NetLapRLS, when integrating multiple networks into a heterogeneous one (Supplementary Information). These results indicated that integrating multiple networks into DTI prediction is not a trivial task, while the network integration procedure of DTINet can simultaneously and effectively capture the underlying topological structures of multiple networks, leading to the improved accuracy of DTI prediction.

### 3.2 DTINet identifies novel drug-target interactions

Next, we analyzed the novel drug-target interactions predicted by DTInet, which have not been reported in existing databases or previous literatures (to the best of our knowledge). The DrugBank database [74] only indicates that clozapine, an antipsychotic drug for treating schizoaffective disorder, can interact with dopamine (DRD) receptors, 5-hydroxytryptamine (HTR) receptors, histamine receptors (HRH), alpha-adrenergic (ADRA) receptors and muscarinic acetylcholine receptors (CHRM1). Our new predictions showed that clozapine can also act on the gamma-aminobutyric acid (GABA) receptors, an essential family of channel proteins that modulate the cognitive functions (Figure 3b. This new prediction can be supported by the previous studies which showed that clozapine can have a direct interaction with the GABA B-subtype (GABA-B) receptors [75] and antagonize the GABA A-subtype (GABA-A) receptors in the cortex [72].

**Figure 3:**
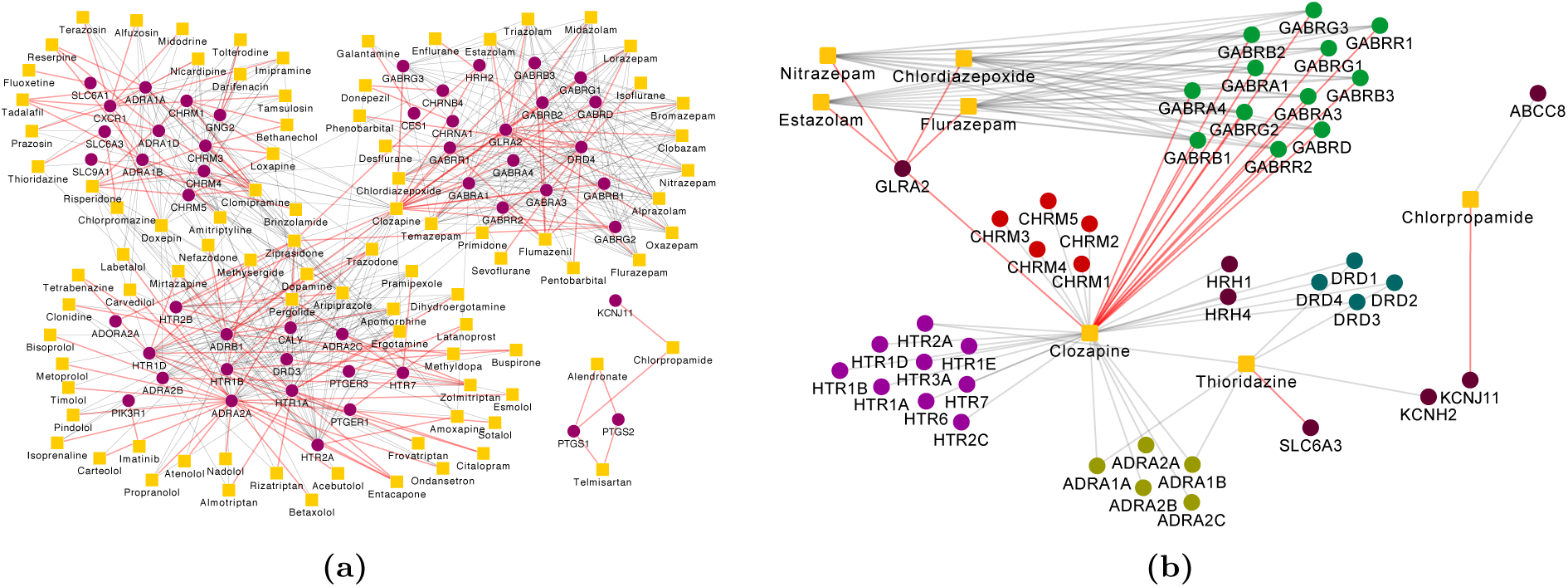
Network visualization of the drug-target interactions predicted by DTINet. (a) Visualization of the overall drug-target interaction network involving the top 150 predictions. Target and drugs are shown in purple circles and yellow boxes, respectively. (b) Network visualization of several examples of novel DTI predictions which can be supported by known experimental or clinical evidence in the literature. The drugs are shown in yellow boxes, while different families of their interacting targets are shown in circles with different colors. In both (a) and (b), known drug-target interactions are marked by grey edges, while the new predicted interactions are shown by red edges.

Chlordiazepoxide, nitrazepam, estazolam and flurazepam are benzodiazepines serving as a class of psychoactive drugs, which are believed to act on the GABA-A receptors [54]. The glycine receptors (GlyRs) are ligand-gated transmembrane ion channel proteins in the central nervous system that play important roles in a wide range of physiological functions, such as the inhibitory synaptic transmission in the brain stem and spinal cord [38]. Our new predictions implied that these benzodiazepine drugs can also interact with the glycine receptors subunit alpha-2 GLRA2 (Figure 3b). This finding agreed with the previous electrophysiological study [58] which indicated that benzodiazepines, such as chlordiazepoxide and nitrazepam, can block the alpha2-containing GlyRs in the embryonic mouse hippocampal neurons through a low-affinity binding manner. Our new predictions along with the previous experimental evidence in [58] may provide important clues for revealing the mechanisms of drug action for benzodiazepines in the central nervous system. In addition, the interaction between clozapine and GLRA2 predicted by DTINet (Figure 3b) was consistent with the previous known experimental results that clozapine is a non-competitive antagonist of GlyRs in the rat hippocampal neurons and thus may contribute to the side-effects associated with the clozapine treatment [37].

Thioridazine is an antipsychotic drug that was widely used to treat schizophrenia and psychosis [59]. The dopamine transporter (also known as the dopamine active transporter, DAT or SLC6A3) is a membrane-spanning protein that mediates the neurotransmission of dopamine from the synapse into neurons and has become an important target for a variety of pharmacologically-active drugs [9]. Our new prediction indicated that there exists an interaction between thioridazine and the dopamine transporter (Figure 3b). Although we did not find direct experimental evidence in the literature to verify this new predicted drug-target interaction, it may be supported from the following two facts. First, thioridazine has been known as a substrate and an inhibitor of the enzyme cytochrome P450 2D6 (CYP2D6) [74, 47]. Second, the enzyme CYP2D6 and the dopamine transporter share a large degree of structural and functional homogeneity [63, 23, 52]. Thus, we can speculate that thioridazine can also act on the dopamine transporter based on these two known evidences.

Chlorpropamide has been known as a first generation sulfonylurea drug that is mainly used to treat the type 2 diabetes mellitus [74]. The subunit Kir6.2 of the ATP-sensitive inward rectifier potassium channel (KCNJ11) is an integral membrane protein that plays important roles in a wide range of physiologic responses, and its mutations are related to many diseases, such as congenital hyperinsulinism [55]. Our prediction showed that chlorpropamide can interact with the KCNJ11 protein (Figure 3b). This predicted drug-target interaction may be supported by the known clinical evidence that sulfonylureas are usually effective drugs for treating most diabetes patients associated with the KCNJ11 mutations [45, 3].

Next, we focused on those novel drug-target interactions among the list of top 150 predictions from DTINet, for which we rarely found known experimental support in the literature. Among the list of these top 150 predictions, most of the new predicted DTIs were relevant (i.e., connected) to the previous known interactions except the interactions between three drugs, including telmisartan, chlorpropamide and alendronate, and the prostaglandin-endoperoxide synthase (PTGS) proteins, which are also called cyclooxygenase (COX) proteins (Figure 3a). COX is a family of enzymes responsible for prostaglandin biosynthesis [64], and mainly includes COX-1 and COX-2 in human, both of which can be inhibited by nonsteroidal anti-inflammatory drugs (NSAIDs) [48]. Apparently, it was difficult to use the correlations between nodes within the DTI network to explain the predicted interactions between these three drugs and the COX proteins. On the other hand, these new DTIs had relatively high prediction scores in the list of the top 150 predictions (Supplementary File 1). In addition, the COX proteins provide a class of important targets in a wide range of inflammatory diseases [40]. Despite the existence of numerous known NSAIDs used as COX inhibitors, many of them are associated with the cardiovascular side-effects [28, 61]. Thus, it is always important to identify alternative COX inhibitors from existing drugs with less side-effects. Given these facts, it would be interesting to see whether the predicted interactions between these three drugs and the COX proteins can be further validated.

Among the aforementioned three drugs, telmisartan has been known as an angiotensin II receptor antagonist that can be used to treat hypertension [21], chlorpropamide has been known as a sulfonylurea drug that acts by increasing insulin to treat type 2 diabetes mellitus [13], and alendronate has been known as a bisphosphonate drug mainly used for treating bone disease, such as osteoporosis and osteogenesis imperfect [4, 16]. Despite our current understanding about the functions of COX-1 and COX-2 proteins and the known indications of telmisartan, chlorpropamide and alendronate, it still remains largely unknown whether these three drugs can also interact with the COX proteins. According to the top 150 predictions by DTINet (Figure 3a and Supplementary File 1), these three drugs can act on the COX proteins. We will further present our validation results on the predicted interactions between these three drugs and COX proteins in the next sections.

### 3.3 Computational docking suggests the binding modes for the predicted drug-target interactions

We preformed both computational docking and experimental assays to validate the new predict interactions between COX proteins and three drugs, including telmisartan, alendronate and chlorpropamide. Here, we mainly present the docking results, and will provide the experimental validation results in the next section. In our structure-based modeling studies, we used the docking program Autodock [42] to infer the possible binding modes of the new predicted interactions between three drugs (i.e., telmisartan, chlorpropamide and alendronate) and the COX proteins.

Our docking results showed that these three drugs were able to dock to the structures of both COX-1 (PDB ID: 3kk6) and COX-2 (PDB ID: 3qmo), and displayed different binding patterns (Figure 4). In particular, all three drugs were fitted into the active sites of both COX-1 and COX-2. More specifically, chlorpropamide displayed similar configurations when binding to COX-1 and COX-2 (Figures 4a and 4b), by forming hydrogen bonds with both residues R120 and Y355, which created a conserved pocket as in those for common NSAIDs [49, 67]. On the other hand, the substitution of V119 in COX-1 by S119 in COX-2 allowed the formation of a different hydrogen bond network in the binding pocket. Moreover, telmisartan and alendronate interacted with residue S530 in addition to residues R120 and Y355 when docked to COX-1 (Figures 4c and 4e), while they were both able to bind to residue S119 when docked to COX-2 (Figures 4d and 4f). Thus, a subtle difference between the binding pockets of those two enzymes may result in different binding modes even for the same drug. These docking results may provide important hints for understanding the structural basis of the predicted drug-target interactions and thus help reveal the underlying molecular mechanisms of drug action.

**Figure 4:**
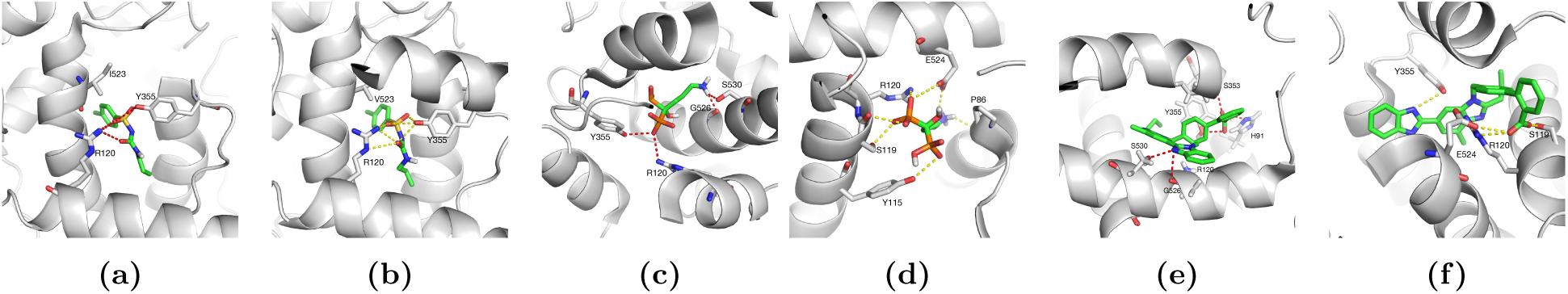
The docked poses for the predicted interactions between three drugs (i.e., chlorpropamide, alendronate and telmisartan) and the COX proteins (i.e., COX-1 and COX-2). (a) Chlorpromamide vs. COX-1; (b) Chlorpromamide vs. COX-2; (c) Alendronnate vs. COX-1; (d) Alendronnate vs. COX-2; (e) Telmisartan vs. COX-1; (f) Telmisartan vs. COX-2. The protein structures of COX-1 and COX-2 were downloaded from the Protein Data Bank (PDB IDs 3kk6 and 3qmo for COX-1 and COX-2, respectively). The structures of the small molecules were obtained from the ZINC [25]. Hydrogen bonds were computed by PyMOL [51] and represented by the red and yellow dashed lines in COX-1 and COX-2, respectively.

### 3.4 Experimental validation of the top-ranked drug-target interactions predicted by DTINet

We further conducted experimental assays to validate the new predicted interactions between the above three drugs, including telmisartan, alendronate and chlorpropamide, and the COX-1 and COX-2 proteins, for which we rarely found other known experimental support in the literature. In the above section, we had performed computational docking to demonstrate that these three drugs can bind to the COX proteins. Here, we further carried out the COX inhibition assays and examined the changes in the gene expressions of the proinflammatory factors after drug treatment (Supplementary Information) to validate these predicted drug-target interactions and demonstrate the new potential applications of these drugs in preventing inflammatory diseases.

We sought to experimentally validate the bioactivities and selectivities of the COX inhibitors predicted by DTINet. First, we tested their inhibitory potencies on the mouse kidney lysates. Similar dose-dependent repression of COX activity was observed for the three drugs (Figures 5a-5c). The IC50 values of telmisartan, alendronate and chlorpropamide for COX activity were measured at 56.14*μ*M 160.2*μ* and 289.5 *μ*M, respectively. The measured IC50 values of the three drugs especially telmisartan were comparable to those of many common NSAIDs, such as celecoxib (COX-1: 82 *μ*M; COX-2: 6.8 *μ*M), ibuprofen (COX-1: 12 *μ*M, COX-2: 80 *μ*M) and rofecoxib (COX-1: >100 *μ*M; COX-2: 25 *μ*M) [26, 27]. Probably alendronate and chlorpropamide were relatively weak inhibitors of COX. It is worth noting that the order of the experimentally measured IC50 values of IC50s of these three drugs was consistent with the ranking of prediction scores in DTINet (Supplementary File 1).

**Figure 5:**
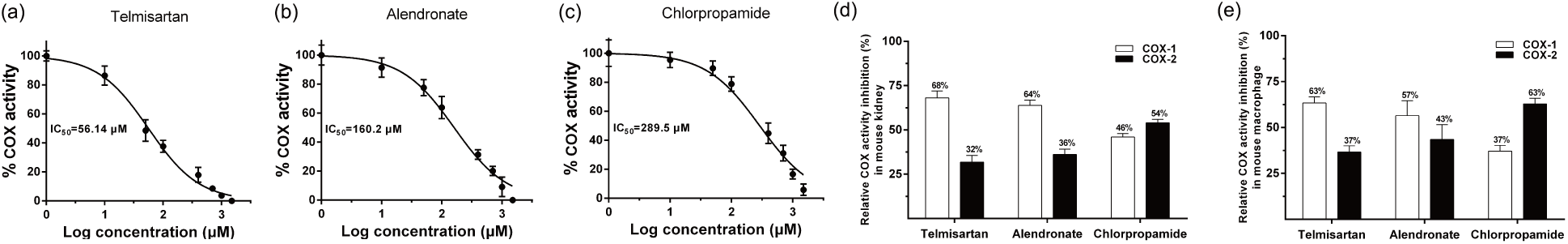
Inhibitory effects of telmisartan, chlorpropamide and alendronate on COX activity in mouse kidney and macrophage lysates. (a)-(c) Inhibition rates of telmisartan, chlorpropamide and alendronate using the mouse kidney lysate. (d) and (e) The relative COX activity inhibition rates of telmisartan, chlorpropamide and alendronate on COX-1 and COX-2, using the tissue extracts from both kidney (d) and macrophage (e) lysates. Here, data show the mean with the standard deviation of three independent experiments, each of which was performed with triplicates.

Next, the tissue extracts from the mouse kidney and the peritoneal macrophages were used for COX selective inhibition assays. Relative inhibition of COX-1 and COX-2 activities was distinguished using SC-560, a potent and selective COX-1 inhibitor, and Dup-697, a potent and time-dependent of COX-2 inhibitor, respectively. Overall, the tests on the tissue extracts from mouse kidney showed that telmisartan and alen dronate preferably inhibited COX-1 (68% and 64%, respectively) over COX-2 (32% and 36%, respectively), while chlorpropamide had a slightly higher inhibition rate on COX-2 (54%) than on COX-1 (46%) (Figure 5d). Similar patterns of COX inhibition selectivity with these drugs were also observed in the peritoneal macrophages (Figure 5e). Overall, the above inhibition assays showed that these three drugs identified by DTINet had a certain level of inhibition affinity and binding selectivity on the family of COX proteins.

The COX inhibitors have been extensively used as nonsteroidal anti-inflammatory drugs (NSAIDs), thus we further tested the effects of the above three drugs on inflammatory responses and thus examined their potential applications in treating inflammatory diseases. Lipopolysaccharide (LPS) was used to stimulate the cultured peritoneal macrophages for the cellular inflammation model. In addition to those three drugs (i.e., telmisartan, chlorpropamide and alendronate) predicted by DTINet, we also considered the potent COX-2 inhibitor Dup-697 and the well-known NSAID ibuprofen for comparison.

A large amount of proinflammatory factors can be generated during the inflammation process [2]. We consequently tested whether the three drugs can suppress the expression of various inflammatory factors in response to LPS stimulation (Figure S7). For TNF-*α* and IL-6, telmisartan exhibited strong inhibitory effect on the LPS induced expression (Figures S7a and S7b). Meanwhile, the induction of the important cytokine IL-1,*β* was also attenuated by each of the three drugs in the peritoneal macrophages (Figure S7c). In particular, telmisartan displayed the strongest suppression effect on IL-1,*β* among all COX inhibitors. For IL-12p35, although both alendronate and telmisartan significantly inhibited its production induced by LPS, telmisartan had much stronger suppression effect than other COX inhibitors (Figure S7d). The LPS-induced production of the immunological defensive factors such as CXCL-1 and iNOS were significantly restrained by the treatment of any of these three drugs (Figures S7e and S7f), which was similar to the results of both Dup-697 and ibuprofen. In summary, these results showed that telmisartan, chlorpropamide and alendronate can reduce the expressions proinflammatory factors in mouse peritoneal macrophages. The observed anti-inflammation effects of these three drugs further extended the above inhibition assay studies and demonstrated their potential applications in preventing inflammatory disease.

Taken together, the above experimental assays validated the novel interactions between the three drugs (i.e., telmisartan, alendronate and chlorpropamide) and the COX proteins predicted by DTINet, which further demonstrated the accuracy of its prediction results and thus provided strong evidence to support its excellent predictive power. In addition, the experimentally validated interactions between these three drugs and the COX proteins can provide great opportunities for drug repositioning, i.e., finding the new functions (i.e., anti-inflammatory effects) of these drugs, and offer new insights into the understanding of their molecular mechanisms of drug action or side-effects of these drugs.

## 4 Discussion

The challenge in network integration mainly stems from the complexity and heterogeneity of datasets. The high-dimensional, incomplete and noisy nature of high-throughput biological data further exacerbates the difficulty. To address this issue, DTINet takes a novel dimensionality reduction technique, which first characterizes the topology of each individual network by applying a network diffusion algorithm (e.g., random walk with restart), and then computes a low-dimensional feature vector representation for each node in the networks to approximate the diffusion information. These low-dimensional feature vectors encode both global and local topological properties for all drug or protein nodes in the networks and are readily incorporable for the downstream predictive models (e.g., matrix completion in this work). We have demonstrated that DTINet can display excellent ability in network integration for accurate DTI prediction and achieve substantial improvement over the previous DTI prediction approaches. Moreover, three novel drug-target pairs predicted by DTINet were also validated by wet-lab experiments, which can provide new insights into the understanding of drug action and drug repositioning.

A future direction of our work is to include more heterogeneous network data in our framework. While we used only four domains (i.e., drugs, proteins, diseases and side-effects) of information in this work, we highlight that DTINet is a scalable framework in that more additional networks can be easily incorporated into the current prediction pipeline. Other biological entities of different types, such as gene expression, pathways, symptoms and Gene Ontology (GO) annotations, can also be integrated into the heterogeneous network for DTI prediction. Although it was only applied to predict missing DTIs in this work, DTINet is a versatile approach and definitely can also be applied to various link prediction problems, e.g., predictions of drug-side-effect associations, drug-drug interactions and protein-disease associations.

## Supplementary Information

### The optimization process of DCA

For simplicity, here we only show the optimization process of DCA for a single input network. The optimization of DCA with multiple networks is a simple extension. To optimize the following objective function of DCA,

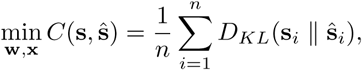

we first express the formula in terms of **w** and **x** based on the definition of KL-divergence and ŝ, that is,

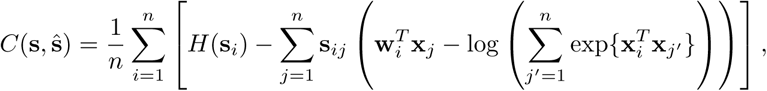

where *H*(⋅) denotes the entropy. Then we compute the gradients of this objective with respect to the parameters **w** and **x**, respectively,

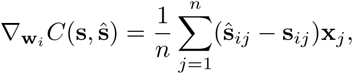

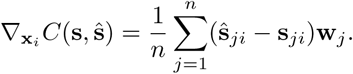

This objective function can be solved using a standard quasi-Newton L-BFGS method to find the lowdimensional vector representations **w** and **x**. Throughout our tests, the vectors **w** and **x** were initialized with uniform random values in [−0.05, 0.05].

As mentioned in the main text, to make the DCA framework more scalable to large biological networks, we use a more efficient matrix factorization based approach, called clusDCA [69], to decompose the diffusion states and obtain their low-dimensional vector representations. Based on the definition of ŝ_*i j*_, we have

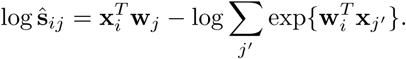

The first term in the above equation corresponds to the low-dimensional approximation of ŝ_*ij*_, and the second term is a normalization factor, which ensures that ŝ_*i*_ is a well defined distribution. We relax the constraint that the entries in ŝ_*i*_ must sum to one by dropping the second term, that is

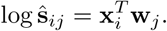

In addition, instead of minimizing the relative entropy between the original and approximated diffusion states, we use the sum of squared errors as the objective function:

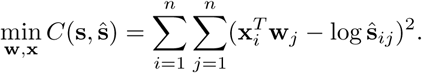

This resulting objective function can be optimized by singular value decomposition (SVD). To avoid taking the logarithm of zero, we add a small positive constant 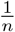 to **s**_*ij*_ and compute the logarithm diffusion state matrix **L** as:

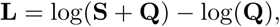

where **Q** ∈ ℝ^*n*×*n*^ with 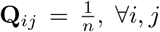 and **S** ∈ ℝ^*n*×*n*^ is the concatenation of **s**_1_,…, **s**_*n*_. With SVD, we decompose **L** into three matrices:

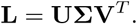

To obtain the low-dimensional vectors **w**_*j*_ and **x**_*i*_ of *d* dimensions, we simply choose the first *d* singular vectors in **U**_*d*_, **V**_*d*_ and the first *d* singular values in Σ_*d*_. More precisely, let **X** = [**x**_1_,…, **x**_*n*_]^*T*^ denote a matrix where each row represents the corresponding low-dimensional feature vector representation of each node in the network, and let **W** = [**w**_1_,…,**w**_*n*_]^*T*^ denote a matrix where each row represents the corresponding vector of the context features. Then, **X** and **W** can be computed as:

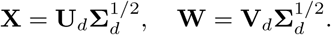

To integrate heterogeneous network data, we extend the above single-network DCA to a multiple-network case. More specifically, let **L** = **{L**^1^,…,**L**^*K*^**}** be the set of logarithm diffusion state matrices based on the set of diffusion states 𝕊 = **{S**^1^,…, **S**^K^**}** from *K* input networks. Then, we optimize the following objective function:

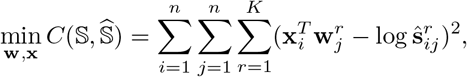

where we assign a network-specific feature **w***_i_^r^* for each node *i* in network *r*, and the node features **x**_*i*_ are shared across all *K* networks. This objective function can also be optimized by SVD.

In the experiments of this work, we use **{x**_*i*_**}** as the low dimensional vector representations of drugs or proteins. Note that **{w**_*i*_**}** can also be used as low dimensional vector representations, but we observed similar performances of these two representations in our local tests.

### Construction of heterogeneous network

A total of four types of nodes and six types of edges, representing diverse drug-related information, were collected from the public databases and used to construct the heterogeneous network for our drug-target interaction (DTI) prediction task.

#### Nodes

We extracted the drug nodes from the DrugBank database (Version 3.0) [32] and the protein nodes from the HPRD database (Release 9) [46]. The disease nodes were obtained from the Comparative Toxicogenomics Database [14]. The side-effect nodes were collected from the SIDER database (Version 2) [34]. In addition, we excluded those isolated nodes, in other words, we only considered those nodes which had at least one edge (see below) in the network.

#### Edges

We imported the known drug-target interactions as well as drug-drug interactions from Drug-Bank (Version 3.0) [32]. The protein-protein interactions were downloaded from the HPRD database (Release 9) [46]. The drug-disease and protein-disease associations were extracted from the Comparative Toxicogenomics Database [14]. We also included the drug-side-effect associations from the SIDER database (Version 2) [34].

**Table S1:** (a) The number of nodes of individual types and (b) the size of individual interaction or association matrices in the constructed heterogeneous network, respectively.

| Type of node | Count | Type of edge | Count |
| --- | --- | --- | --- |
| Drug | 708 | Drug-Protein | 1,923 |
| Protein | 1,512 | Drug-Drug | 10,036 |
| Disease | 5,603 | Drug-Disease | 199,214 |
| Side-effect | 4,192 | Drug-Side-effect | 80,164 |
| Total | 12,015 | Protein-Protein | 7,363 |
|  |  | Protein-Disease | 1,596,745 |
|  |  | Total | 1,895,445 |

For the input homogeneous interaction networks (e.g., drug-drug interaction network), we compute the “diffusion state” of each drug or target by directly running the RWR algorithm on each of these networks. For the association networks, i.e., drug-side-effect, drug-disease, and protein-disease association networks, we construct the corresponding similarity networks based on the Jaccard similarity coefficient and then run the RWR process on these similarity networks. Jaccard similarity is a common statistic used to characterize the similarity between two sets of objects. Taking the drug-side-effect association network as an example, we use the following formula to measure the similarity between drug *i* and drug *j*:

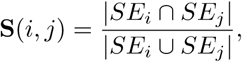

where *SE_i_* denotes the set of side-effects of drug *i*. Then we run the RWR procedure on this similarity network to obtain the diffusion states of drugs. In the same manner, we can construct the similarity networks of proteins.

In addition to the above interaction or association-based similarity networks, we construct a drug similarity network based on the chemical structures of drugs, in which the similarity score between a pair of two drugs is calculated using the Tanimoto coefficient [22] using the product-graphs of their chemical structures. We also construct a protein similarity network based on genome sequences, in which the similarity score between a pair of two proteins is computed using the Smith-Waterman score [56] based on their primary sequences.

Overall, we construct four similarity networks for drugs, based on (i) drug-drug interactions, (ii) drug-disease associations, (iii) drug-side-effect associations, and (iv) chemical structures. Similarly, we construct three similarity networks for proteins, based on (i) protein-protein interactions, (ii) protein-disease associations, and (iii) genome sequences. With these similarity networks, we can learn the low-dimensional feature vector representations of drugs and proteins, by first performing diffusion separately on individual networks and then jointly optimizing the feature vectors under the compact feature learning framework.

### Parameters setting of DTINet

For DTINet, we observed stable performance of DTINet for different values of the restart probability *p_r_* between 0.5 and 0.8 (Figure S6a). For all the test results in this work, the restart probability *p_r_* was set to 0.5. After the compact feature learning, we obtained an *f_d_*-dimensional vector for each drug and an *f_t_*-dimensional vectors for each target. We observed robust results over a wide range of choices for the *f_d_* and *f_t_* parameters (Figure S5). In our experiments, we set *f_d_* = 100 and *f_t_*= 400, which were equal to 10-20% of the dimensionality of the original vectors describing the diffusion states. We also evaluated the prediction performance of DTINet with respect to different choices of the latent dimensionality parameter *f_k_* and observed stable performance of DTINet over a wide range values of this parameter (Figure S6b). In all experiments of this work, we set the value of the latent dimensionality parameter to *f_k_* = 50, which was roughly equal to 50% of the dimension of the feature vectors of drugs, or 10% of the dimension of the feature vectors of proteins.

### Baseline methods

We compare our method against four previously-proposed methods, including the bipartite local models, the Laplacian regularized least square, the heterogeneous network model and the collaborative matrix factorization. We briefly describe these methods below.

1. Bipartite local model with neighbor-based interaction-profile inferring (BLMNII) [39]: This method is a combination of the bipartite local model (BLM) and the neighbor-based interaction-profile inferring (NII). The BLM framswork models the drug-target interaction prediction task as a binary classification problem in a bipartite graph. Suppose that we want to predict whether drug *d*_i_ interacts with target *t_j_*. The BLM method first focuses on drug *d_i_* and assigns a label +1 to all the known targets that interact with drug *d*_i_, and −1 otherwise. Then BLM uses the protein similarity matrix as a kernel matrix to train a support vector machine (SVM). Such a process is also performed in a reverse way, that is, BLM also labels each known drug by whether it interacts with target *t_j_* or not, and then trains an SVM based on the drug similarity matrix. The final prediction of whether drug d_i_ interacts with target *t_j_* is then derived based on the average prediction score from both directions. The NII procedure incorporates the neighbors interaction profiles into the BLM method to train the model and enable the prediction for new drugs or targets.
2. Laplacian regularized least square (NetLapRLS) [76]: This method employs a semi-supervised learning algorithm, i.e., Laplacian regularized least square, for DTI prediction, which utilizes available labeled data of DTI pairs and incorporates similarity and interaction kernels to improve the prediction. NetLapRLS attempts to estimate the interaction scores **F**_d_ and **F**_*t*_ based on the drug and protein domains, respectively. For example, the interaction scores **F**_*d*_ are obtained by minimizing the squared loss between the known DTI matrix **P** and **F**_*d*_ with a regularized term of **F**_*d*_ and **S**_*d*_, where **S**_*d*_ is the similarity network of drugs. The final prediction **F** is obtained by averaging the results derived from both **F**_*d*_ and **F**_*t*_.
3. Heterogeneous network model (HNM) [70]: This method builds a three-layer heterogeneous network consisting of three types of nodes: drug, target and disease nodes. Then it iteratively propagates interaction or association information in the heterogenous network using random walk with restart. The iterative updating rule is given by

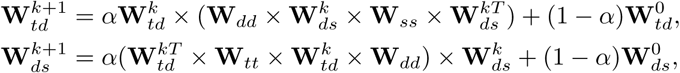

where **W***_td_^k^* and **W***_ds_^k^* stand for the weights on the target-drug and drug-disease association links in the *k*th iteration, respectively; **W***_td_^0^* and **W***_ds_^0^* represent the target-drug and drug-disease association matrices defined by the input data; **W**_*dd*_ stores both drug interaction and similarity information, which is basically computed from the averaging result derived from both of the drug-drug interaction and drug-drug similarity matrices; **W**_*ss*_ represents the disease-disease similarity matrix; and **W**_*tt*_ represents the protein-protein interaction matrix derived from the input data. The final DTI prediction scores are obtained from matrix **W**_*td*_ after convergence.
4. Collaborative Matrix factorization (CMF) [79]: This method learns the feature vector matrices **X** and **Y** for drugs and targets, respectively, by minimize the following objective function:

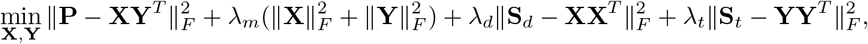

where *S_d_* and *S_t_* represent the drug and target similarity matrices, respectively, and *λ_m_*, *λ_d_* and *λ_t_* represent the regularization coefficients.

Note that HNM and CMF are designed to integrate heterogeneous information, while BLMNII and NetLapRLS mainly focus on solving the DTI prediction problem on a single network. To make a fair comparison, we implemented an extended version for both BLMNII and NetLapRLS, in which we summarized our heterogeneous network (Figure 2a) into a single network for both BLMNII and NetLapRLS, using the following integration process [68]. In particular, we combined multiple networks into a single network by assigning the edge weight 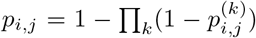, where 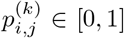 is the interaction probability or similarity between node *i* and node *j* in network *k* ∈ {1, 2,…, *K*}, where *K* stands for the total number of networks.

### Experimental validation

#### Reagents

LPS (L-2360) and 4% sterile thioglycollate were purchased from Sigma-Aldrich (St. Louis, MO, USA). IFN-*γ*(315-05) was purchased from PeproTech (New York, USA). Chlorpropamide (S4166), telmisartan (S1738), alendronate (S1624) and ibuprofen (S1638) were purchased from Selleck Chemicals (Houston, TX, USA). COX Fluorescent Activity Assay Kit (700200) was purchased from Cayman Chemical Company (Ann Arbor, MI, USA).

#### Cell culture

C57BL/6J mice (10 weeks old) were obtained from Vital River (Beijing, China) and were housed under controlled temperature (22 °C ± 2 °C) and humidity (40–60%) with a 12 h light/dark cycle. Mice were injected intraperitoneally with 1 ml of 4% sterile thioglycollate and sacrificed 3 days later. Peritoneal macrophages were isolated from peritoneum by lavage using 20 ml DMEM and seeded into 6 well plates using one hundred million cells/well in DMEM of 10% FBS. Non-adherent cells were removed 6 h later, whereas adherent cells were refed with DMEM of 10% FBS and allowed to recover overnight. Macrophages were treated with chlorpropamide, telmisartan and alendronate for 24h and then pre-incubated with DMEM-10% FBS for 2h before treatment of LPS (10 ng/ml). Macrophages were pre-treated with DuP-697 and SC-560 for 12 h before treatment with telmisartan, alendronate and chlorpropamide and then incubated with IFN-*γ* (10 ng/ml) for 12 h following LPS stimulation (10 ng/ml) for 6 h. The concentrations of telmisartan, alendronate and chlorpropamide treatment were determined based on previous research [36, 62, 19], while those of the chemical probe Dub-697 and the known NSAID ibuprofen were determined according to the indications of the assay kit and previous binding studies in the literature [1, 50, 57], respectively. Cells were harvested for subsequent analysis.

#### COX fluorescent activity

Following stimulation, kidneys were harvested from mice, and macrophages from the above treatment were homogenized in 5 ml of cold PBS containing protease inhibitors and centrifuged at 10,000 g for 15 minutes at 4 °C. The supernatant was assayed by the COX fluorescent activity assay kit according to the manufacturer’s instructions.

#### Real-time PCR analysis

Total RNA was extracted from the whole-cell lysates using the Trnzol-A+ reagent (Tiangen, Cat. no. DP421, China). Reverse transcription was performed using TIANScript RT Kit (Tiangen, Cat. no. KR104-02, China). All real-time PCR reactions were carried out on ABI ViiA^TM^ 7 Real-Time System (Life Technologies, USA) using TransStart Top Green qPCR SuperMix (Transgen, Cat. no. AQ131-03, China). The formula 2^−ΔΔCt^ was used to calculate the relative expression. The expression of the housekeeping gene GAPDH was used as an internal control.

### Top 150 predictions of drug-target interactions by DTINet

#### Supplementary File 1

A list of top 150 predictions of drug-target interactions by DTINet is available at http://github.com/luoyunan/DTINet.

## Acknowledgments

This work was supported in part by the National Basic Research Program of China (Grant 2011CBA00300 and 2011CBA00301), the National Natural Science Foundation of China (Grant 61033001, 61361136003, 61472205 and 81470839), the China’s Youth 1000-Talent Program, the Beijing Advanced Innovation Center for Structural Biology, and the Tsinghua University Initiative Scientific Research Program (Grant. 20161080086). J.P. received support as an Alfred P. Sloan Research Fellow. We acknowledge the support of NVIDIA Corporation with the donation of the Titan X GPU used for this research.

## Author contributions

Y.L., J.P., L.C. and J.Zeng conceived the research project. J.Z. supervised the research project. Y.L., J.P. and J.Zeng designed the computational pipeline. Y.L. implemented DTINet and performed the model training and prediction validation tasks. X.Z. and L.C. performed the experimental validation task and analyzed the validation results. J. Zhou carried out the computational docking and data analysis tasks. Y.L., X.Z., J.Zhou, J.Y., Y.Z., W.K., L.C. and J.Zeng analyzed the novel prediction results. Y.L. and J.Zeng wrote the manuscript with support from all authors.

---

**Figure S1:**
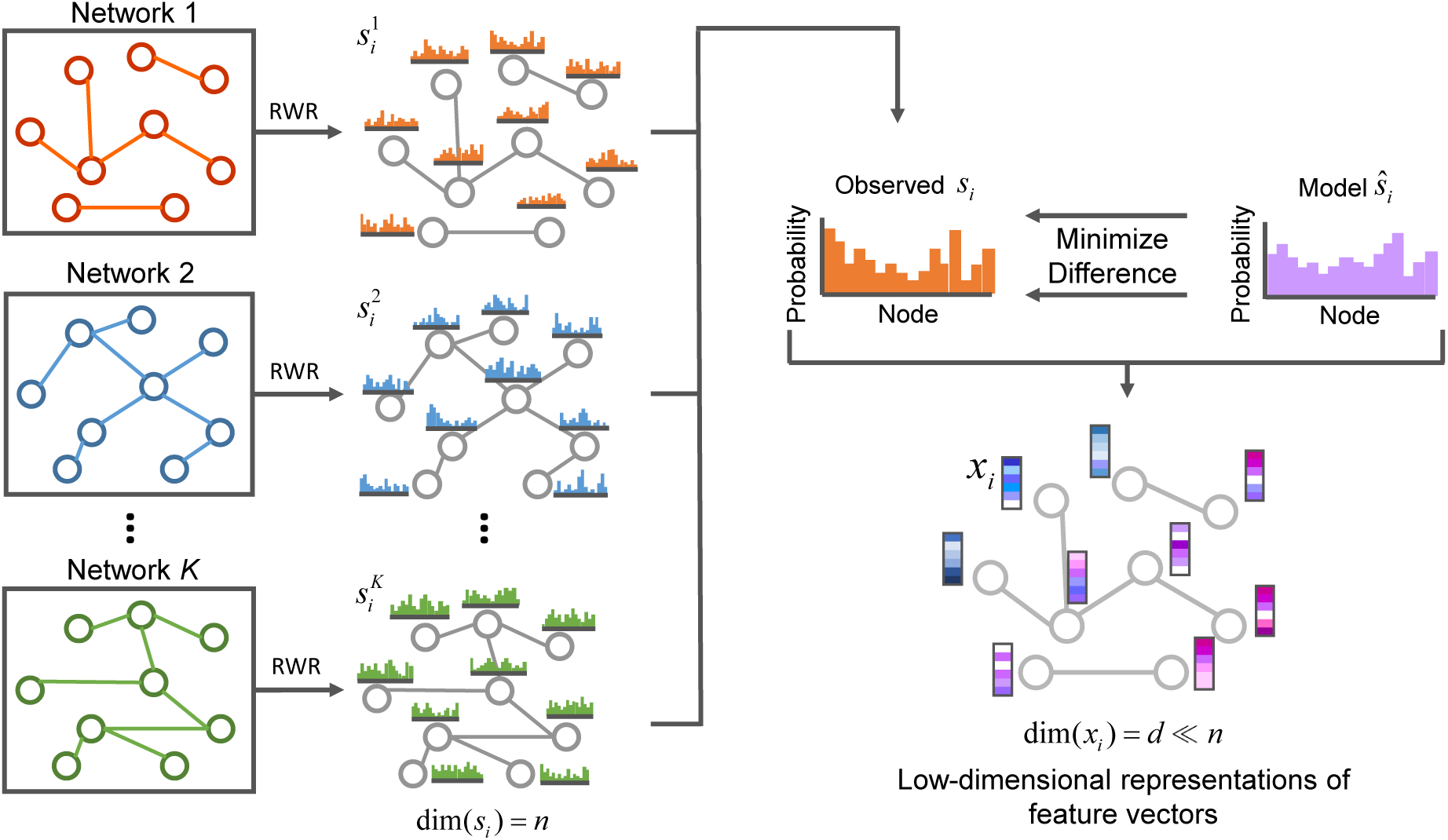
Schematic illustration of compact feature learning. The random walk with restart (RWR) algorithm is first used to compute the diffusion states of individual networks. Then the low-dimensional representations of feature vectors for individual nodes are obtained by minimizing the difference between the diffusion states *s*_i_ and the parameterized multinomial logistic models *ŝ_i_*. The learned low-dimensional feature vectors encode the relational properties (e.g., similarity), association information and topological context of each node in the heterogeneous network. More details can be found in the main text.

**Figure S2:**
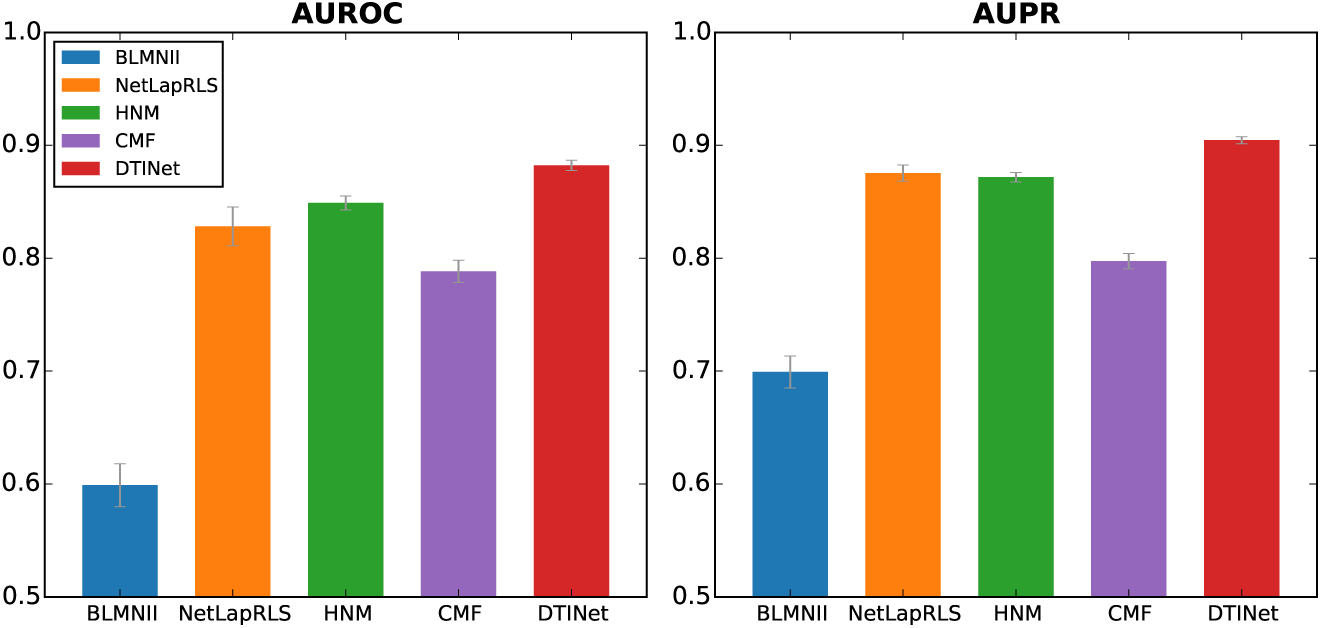
DTINet outperforms other state-of-the-art methods for DTI prediction on a dataset in which homologous proteins were excluded. We performed a ten-fold cross-validation procedure to compare the prediction performance of DTINet to that of four state-of-the-art DTI prediction methods, i.e., HNM, CMF, and the extended versions of BLMNII and NetLapRLS. Performance of each method was assessed by both the area under ROC curve (AUROC) and the area under precision-recall curve (AUPRC). All results were summarized over 10 trials. A pair of two proteins are said to be homologous if their sequence identity score is above 40%.

**Figure S3:**
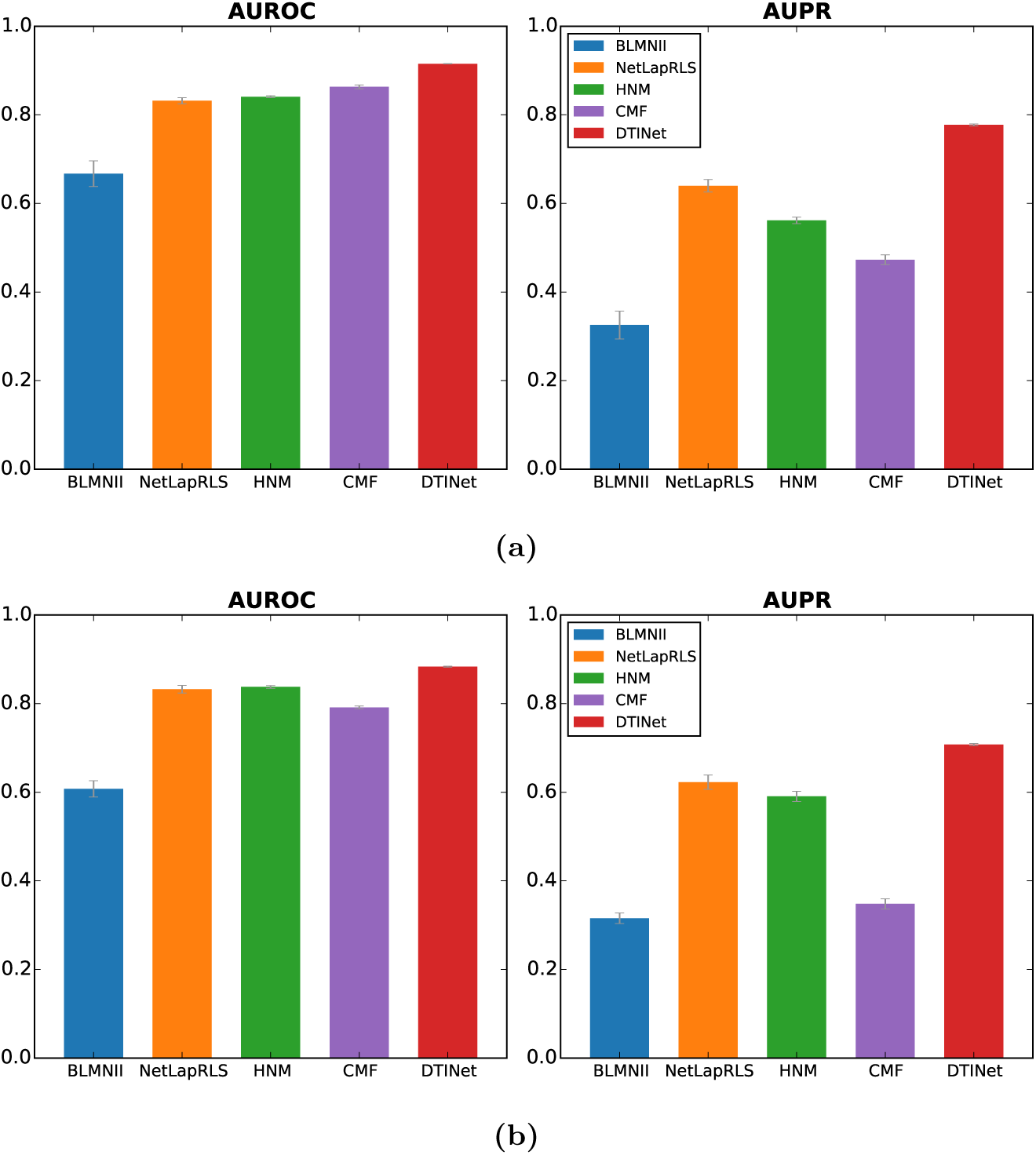
Performance comparison between DTINet and other state-of-the-art methods on a skewed dataset. The number of randomly chosen non-interacting drug-target pairs (i.e., negative samples) was 10 times more than the number of known interacting drug-target pairs (i.e., positive samples). All results were summarized over 10 trials of ten-fold cross-validation. (a) All methods were trained and tested on the original collected dataset (see the main text), without removing any homologous proteins. (b) All methods were trained and tested on a modified dataset, in which homologous proteins were excluded. The cutoff of sequence similarity score 40% was used to define a pair of homologous proteins.

**Figure S4:**
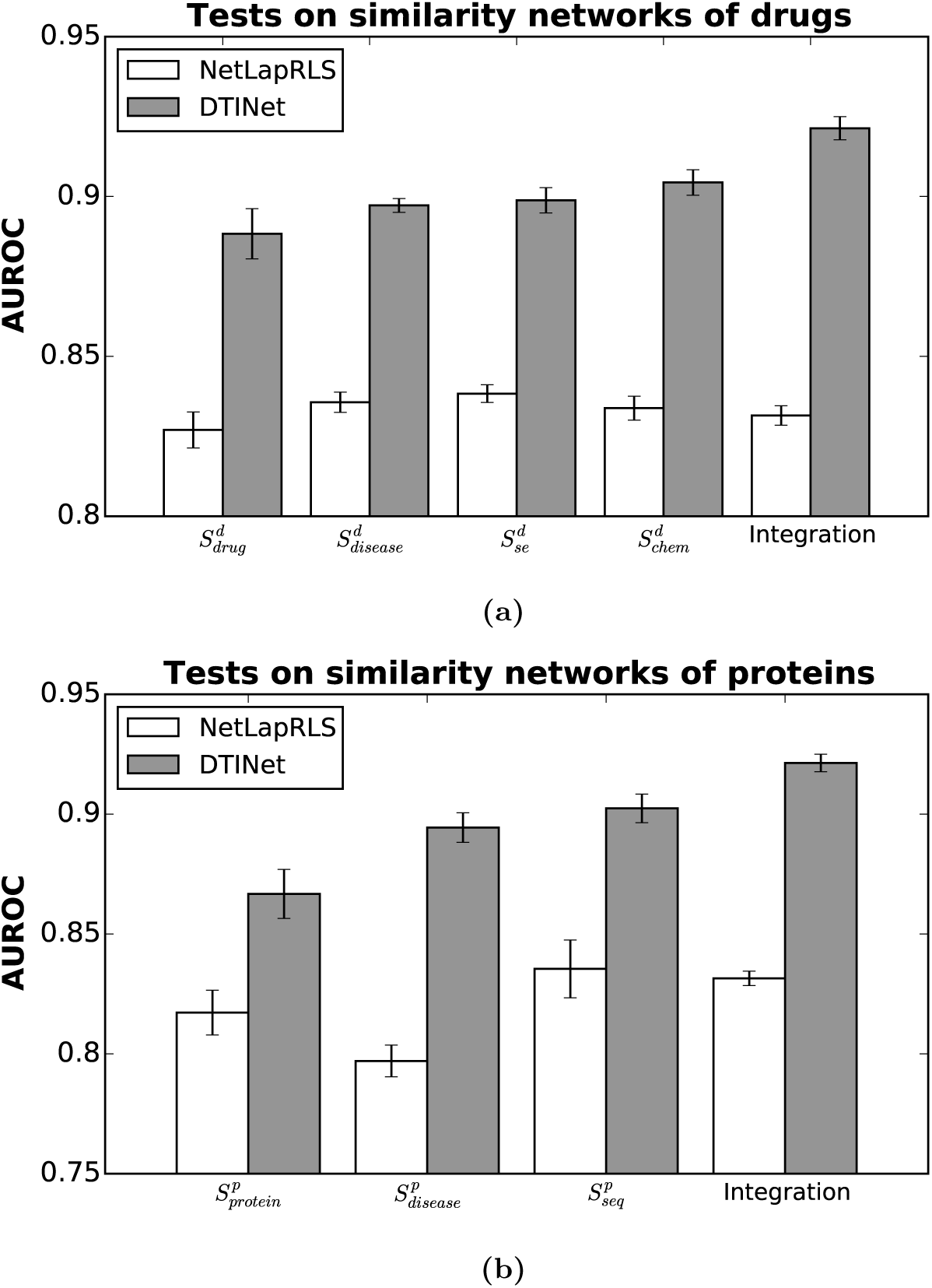
A comparative study on the prediction performance of DTINet and NetLapRLS on individual networks and their integration. (a) The test results on individual similarity networks of drugs and their integration, where *S_drug_^d^*, *S_disease_^d^*, *S_se_^d^* and *S_chem_^d^* represent the similarity networks in which the similarity score between a pair of drug nodes was computed based on the profiles of drug-drug interactions, drug-disease associations, drug-side-effect associations and chemical structures, respectively. (b) Tests on individual similarity networks of proteins and their integration, where *S_protein_^p^*, *S_disease_^p^* and *S_seq_^p^* represent the similarity networks in which the similarity score between a pair of protein nodes was computed based on the profiles of protein-protein interactions, protein-disease associations and primary sequences, respectively. An extended version of NetLapRLS (see Supplementary Information text) was used to combine all similarity networks to perform DTI prediction. All results were summarized over 10 trials of ten-fold cross-validation.

**Figure S5:**
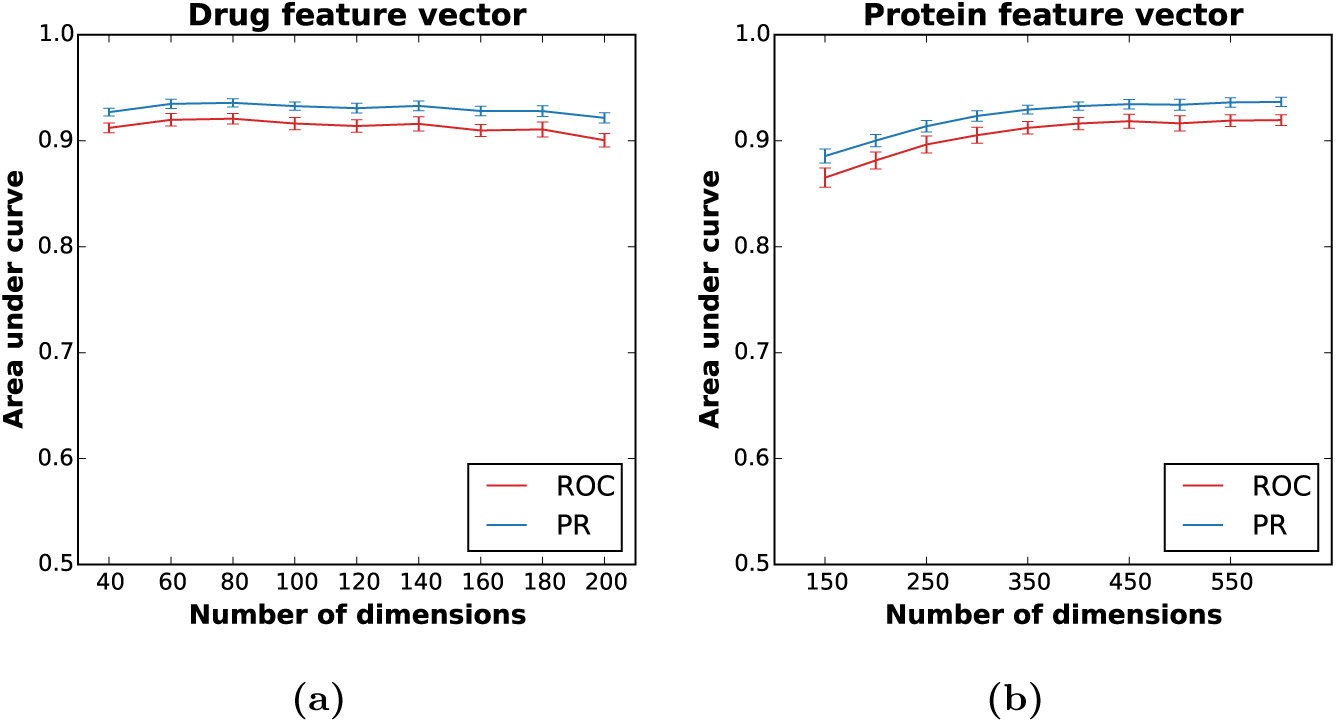
Robustness of DTINet with respect to the number of dimensions of feature vectors. We evaluated the sensitivity of the prediction performance of DTINet with respect to different numbers of dimensions of the feature vectors of drugs (a) and proteins (b). We tested the dimensions of the feature vectors of drugs (*f_d_*) and proteins (*f_t_*) in a range that are roughly equal to 10%-30% of the dimensionality of the original vectors describing the diffusion states. DTINet had stable prediction performance over a wide range the dimensions of the feature vectors. Prediction performance was evaluated in terms of both the area under the receiver operating characteristic curve (ROC) and the area under the precision recall (PR) curve. All results were summarized over 10 trials of ten-fold cross-validation.

**Figure S6:**
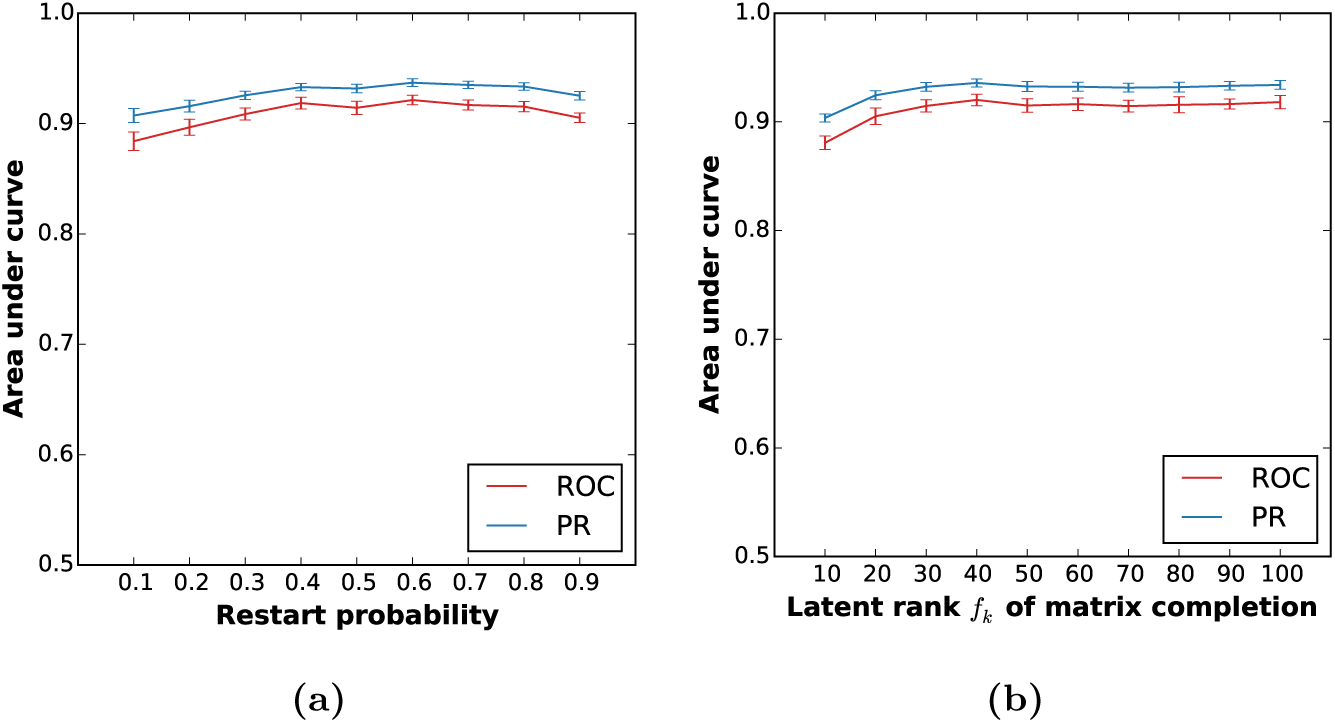
Robustness of the prediction performance of DTINet with respect to the restart probability and the latent rank of matrix completion. We tested the prediction performance of DTINet with respect to different values of the restart probability (a) and the latent rank of matrix completion (b). DTINet was robust to different choices of the restart probability and the latent rank parameter. Prediction performance was evaluated in terms of both the area under the receiver operating characteristic curve (ROC) and the area under the precision recall (PR) curve. All results were summarized over 10 trials of ten-fold cross-validation.

**Figure S7:**
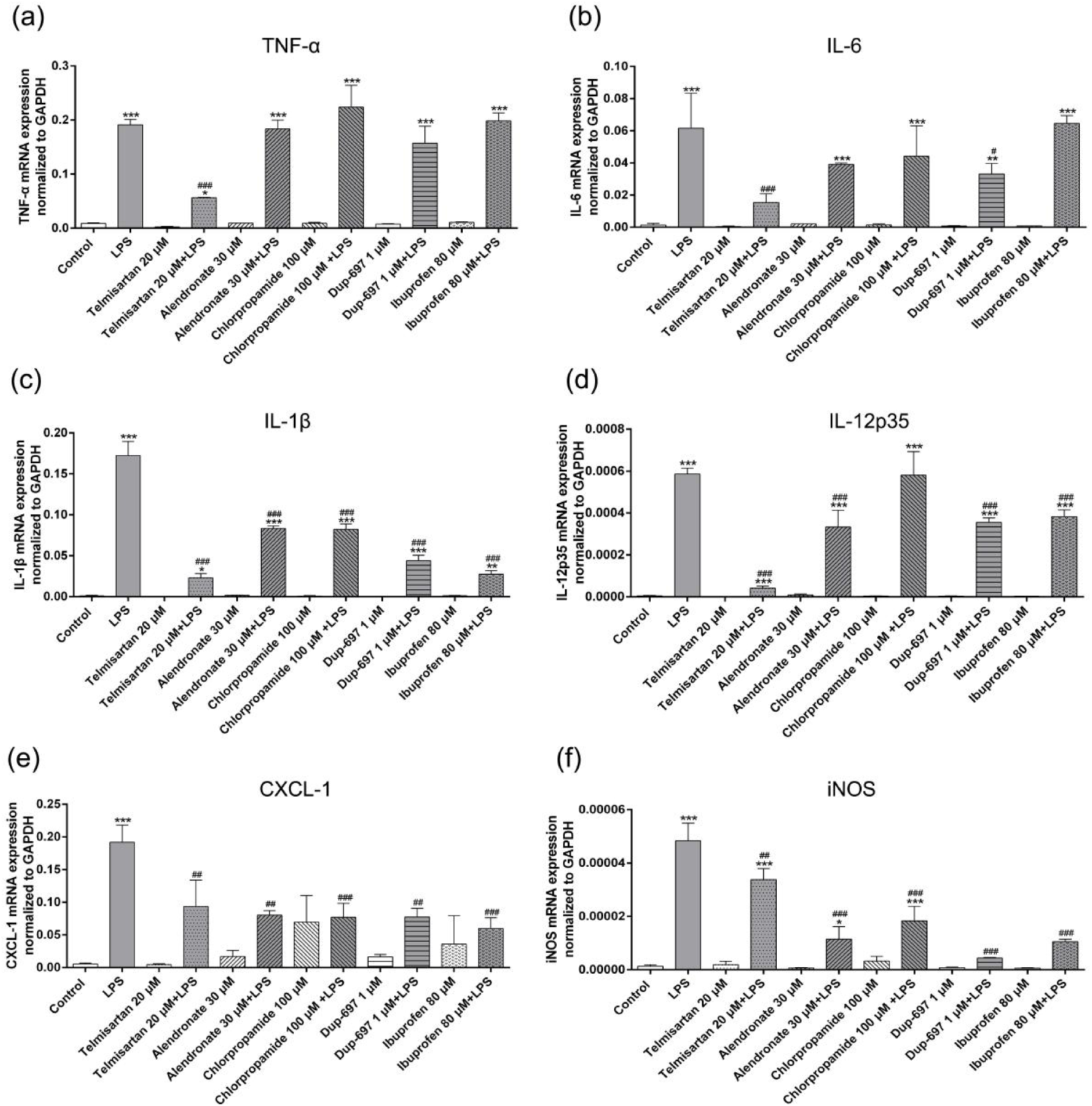
The real-time PCR (RT-PCR) analysis of the proinflammatory factors on the LPS-stimulated macrophages. (a)–(f) The RT-PCR analysis of mRNA expressions of TNF*α*, IL-6, IL-1,*β*, IL-12p35, CXCL-1 and iNOS normalized relative to that of GAPDH, respectively. Control, macrophages without LPS treatment. *: *P <* 0.05; **: *P* < 0.01; ***: *P* < 0.001, compared to the samples without LPS treatment. ##: *P* < 0.01; ###: *P* < 0.001, compared to the samples treated with LPS. n=3. Tukey’s multiple comparison test was used. Here, data show the mean with the standard deviation of three independent experiments, each of which was performed with triplicates. The concentrations of the COX inhibitors were determined according to the indications of the assay kits and the previous binding studies in the literature (see Supplementary Information text).

## Footnotes

This paper was selected for oral presentation at RECOMB 2017 and an abstract is published in the conference proceedings.

